# Vocal efficiency in crows

**DOI:** 10.1101/2025.07.03.663010

**Authors:** Claudia A.F. Wascher, Mason Youngblood

## Abstract

Many communicative systems have been selected for efficiency, shaped by the trade-off between information transmission and energetic or temporal constraints. Linguistic laws such as Menzerath’s law—predicting shorter elements in longer sequences—have emerged as widespread principles across vocal communication in many species. While these laws have been predominantly studied at the species level, the influence of individual and social factors remain underexplored. In this study, we investigated adherence to Menzerath’s law in the vocal communication of carrion crows, *Corvus corone corone*, hooded crows, *Corvus corone cornix* and hybrids. Our findings show that crow call sequences adhere to Menzerath’s law, with shorter calls occurring in longer sequences, demonstrating structural efficiency in vocal communication. In carrion crows specifically, we analysed call sequences in relation to individual characteristics (sex, age) and social variables (group size, dominance status, strength of affiliative relationships). Interestingly, adherence to Menzerath’s law was stronger in males and younger individuals, while no effects were found for group size, dominance, or affiliative relationships. This study provides the first evidence of Menzerath’s law in corvid vocal communication and suggests that individual-level traits, rather than broader social dynamics, may shape vocal efficiency. These findings broaden our understanding of widespread principles in animal communication and raise new questions about the ontogeny and flexibility of vocal efficiency in complex social species.

## Introduction

Biological systems have evolved to operate efficiently under the constraints of energy and time, ensuring survival and reproductive success (Igamberdiev, 2023). This is also the case for a wide range of communication systems, such as human language (Gibson et al., 2019) and vocal communication in non-human animals (Endler, 1993; Semple et al., 2022). Communication systems typically evolve to convey critical information accurately, which often requires an increase of signal complexity. For example in the context of mating, increased signal complexity is associated with increased mating success (Choi et al., 2022; Pollard & Blumstein, 2012). In an anti-predator context, complex signals can convey information such as predator type and level of risk (Suzuki, 2014) and social factors such as increased group sizes, and cohesiveness of social bonds requires increasingly complex communication systems such as increases in vocal repertoires or diversity of signals (Peckre et al., 2019). The need for complexity for information transmission needs to be balanced with time and energy constraints, consequently selecting communication systems for efficiency and allowing to optimize the benefits resulting from information transmission while reducing the costs of signal production (Endler, 1993).

One widespread way to reduce costs of communicating an increase signalling efficiency is the reduction of communication time, for example by minimizing the length of a signal. In the context of human language, this efficiency is traditionally quantified as linguistic laws, such as Zipf’s law of brevity, which predicts a negative relationship between the length of words and how often they are used (Zipf, 1949) and Menzerath’s law, which predicts that longer sequences will be comprised of shorter items to balance production costs (Menzerath, 1954). In recent years, empirical evidence is accumulating suggesting that these information theoretical principles are widespread across a wide range of taxa and communication modalities. Menzerath’s law is common for example in bird song (Favaro et al., 2020; James et al., 2021; Lewis et al., 2023; Youngblood, 2024), primates (Gustison et al., 2016; Huang et al., 2020; Valente et al., 2021), bats (Zhang et al., 2024), or whale vocalisations (Youngblood, 2025). Adherence to Menzerath’s law is not limited to vocal communication but can also be seen in gestural communication (Heesen et al., 2019; Safryghin et al., 2022).

While adherence to linguistic laws seem widespread in non-human animal vocal communication systems, studying social and ecological factors shaping the phenomenon is of interest to further understand the evolutionary pressures shaping efficiency in animal communication systems. Marmoset monkeys, *Callithrix jacchus*, can be trained in a vocal conditioning experiment adopt several vocal compression strategies such as increasing call rate, decreasing call duration and increasing the proportion of short calls, which are consistent with the principles of Zipf’s law of brevity (Risueno-Segovia et al., 2023). However, previous studies on common marmoset and the golden-backed uakari, *Cacajao melanocephalus*, fail to find empirical support for Zipf’s law in vocalisations (Bezerra et al., 2011). In rock hyraxes, *Procavia capensis*, Zipf’s law could only be shown in male but not female calls (Demartsev et al., 2019). In different species Menzerath’s Law is only present in certain call-types (tarsiers monkeys, *Tarsius spectrumgurskyae*, titi monkeys, *Plecturocebus cupreus* and gibbons, *Plecturocebus cupreus:* Clink & Lau, 2020; Hainan frilled treefrogs, *Kurixalus hainanus:* Deng et al., 2024).

In the present study we investigated adherence to Menzerath’s law in two subspecies of carrion crows, carrion crows, *Corvus corone corone*, hooded crows, *Corvis corone cornix* and hybrids (thereafter crows). Crows belong to the family of corvids, a large family of birds belonging to suborder of oscine passerine birds, with vocal communication playing an important role in the socio-ecology of corvids (Wascher & Reynolds, 2025). Corvids mostly produce calls, i.e., short, distinct vocalisations and not song, i.e., heterogeneous, combinatory vocalisations consisting of notes or phrases that are arranged in a specific order and often repeated (Sandoval & Graham, 2025). Previously, Menzerath’s law was mostly investigated in song, not calls. Further, Menzerath’s law is mostly investigated on species level and whether it is present in a certain species or not and only recently individual level effects have been described in Java sparrows, *Padda oryzivora* (Lewis et al., 2023). In the present study, we are aiming to investigate how individual factors, such as sex and age as well as social factors like group size, dominance and affiliative relationship status affect adherence to Menzerath’s law. Corvids show high levels of flexibility in their vocalisations, for example common ravens, *Corvus corax*, flexibly adjust calls to audience composition (Szipl et al., 2018) and show similarity in long-distance calls between pair-partners (Luef et al., 2017). Carrion crows have volitional control over calls (Brecht et al., 2019) and can flexibly produce a variable number of vocalisations, with the acoustic features of the first vocalisation being predictive of the total number of calls in a sequence (Liao et al., 2024). Corvids are open-end vocal learners, which means they can acquire new vocalizations throughout their lifetime and not only in a specific sensitive period (Brenowitz et al., 1997). Many corvid species are mimics, allowing them to copy sounds from the environment or conspecifics (Wascher et al., 2025). As vocal communication plays an important role in allowing corvids to navigate social and ecological challenges, we expect widespread rules of signalling efficiency to apply to their calls. We focus on the Menzerath-Altmann law—a precise and more robust mathematical form of Menzerath’s law (Altmann, 1980; Torre et al., 2021). Data was collected from carrion and hooded crows as well as hybrids, although we mostly focus on carrion crows as sample size for hooded crows and hybrids is very small (two individuals each).

In carrion crows, we further investigate whether within the species adherence to Menzerath’s law varies, depending on individual factors, such as sex and age and social factors (group size, dominance status and strength of affiliative relationships). We expect weaker adherence to Menzerath’s law in younger individuals compared to adults, as they are expected to still be learning vocalisations patterns. As vocal communication is important for both male and female crows, we do not have specific predictions regarding a sex-based difference how strongly Menzerath’s law is observed in carrion crows. Group size is often considered a driving factor of ‘social complexity’ and associated with this ‘vocal complexity’ (Freeberg & Krams, 2015; Krams et al., 2012; Manser et al., 2014). We do expect a higher need for efficiency in larger groups to allow for coordinated vocalisations and behaviour, hence adherence to Menzerath’s law is expected to be higher in larger groups. Lastly, we also investigate effects of dominance and strength of affiliative relationships. We expect the ability to communicate efficiently and hence adherence to Menzerath’s law to be positively correlated with dominance status and strength of affiliative relationships in the group.

## Methods

### Ethical note

The current study was conducted in two populations of captive crows, housed in large outdoor enclosures in Navafría, Castilla y León, Spain (42°36’33 N 5°26’56) and Grünau, Upper Austria (47°51’02 N 13°57’44 E). Crows remained in captivity prior to, during and remained after the presented study. Aviary sizes varied between populations and groups from 20 to 72 m^2^ and are always equipped with wooden perches, natural vegetation and rocks. An enriched diet consisting of fruit, vegetables, bread, meat and milk products is provided daily. In both locations water is available *ad libitum* for both drinking and bathing. Keeping of captive birds in Spain was authorized by Junta de Castilla y León (núcleo zoologicó 005074) and in Austria under a licence issued to the Cumberland game park Grünau (AT00009917).

### Study subjects

Data for the present study has been collected from 2010 to 2015. As data for the present study has been taken over the extended period time, group composition and study subjects changed in different phases, which is summed up in table 1. Most study subjects are carrion crows, however in the Austrian population there were also two hooded crows and two hybrids (table 1). Since a recent taxonomic update, carrion crows and hooded crows are considered subspecies of the same species (AviList Core Team, 2025). Birds were kept in different group compositions, which reflect natural conditions in the wild: (A) ‘flock’ or ‘family’: groups of three and more individuals, mostly juvenile individuals grouped together but in one case a pair successfully reproducing; (B) ‘pair’: Adult individuals are mostly kept in male-female pairs; in some cases also in trios and due to death of partner for a limited amount of time as singles. All birds were hatched in the wild and most were hand-raised and brought into captivity before fledging, except Valencia and Xufa, who hatched in captivity. Groups were visually but not acoustically separated from each other. For more information on study subjects a see Wascher (2021) and Wascher et al. (2019). As an estimate for strength of affiliative relationships for each subject we calculated a Composite Sociality Index (CSI) and as an estimate for dominance rank, we calculated an Elo-rating (ELO) for each subject. We recorded a total of 2,079 individual focal observations (899 in the Austrian population; 1,180 in the Spanish population) lasting five minutes each. All occurring behaviours were recorded but for this study, we focused on the frequencies of agonistic behaviour (threat, chase flight and fight) and affiliative behaviours (allopreen and contact sit). We recorded the identity, role (initiator/receiver) of interacting individuals and the outcome of the agonistic interaction (winner/loser), with the loser of an agonistic interaction defined as the individual that retreated. CSI was calculated according to (Silk et al., 2010). Following Archie et al. (2014), we corrected for a different observational effort between individuals by regressing CSI against the number of observations per individual. We calculated a mean CSI for each individual across all possible dyads, representing the average strength of affiliative relationships. The relative success levels of individuals in agonistic encounters was calculated as an Elo-rating in the R package ‘aniDom’ (version 0.1.4; Sánchez-Tójar et al., 2018). Each individual was rated based on the outcome of each discrete interaction (winner/loser) and the (predicted) probability of that outcome occurring (Neumann et al., 2011). Similar to above, we corrected for a different observational effort between individuals by regressing Elo-rating against the number of observations per individual. Details of behavioural data collection and calculation of CSI and Elo-rating are described in Wascher (2021) and Wascher et al. (2019).

**Table 1.**
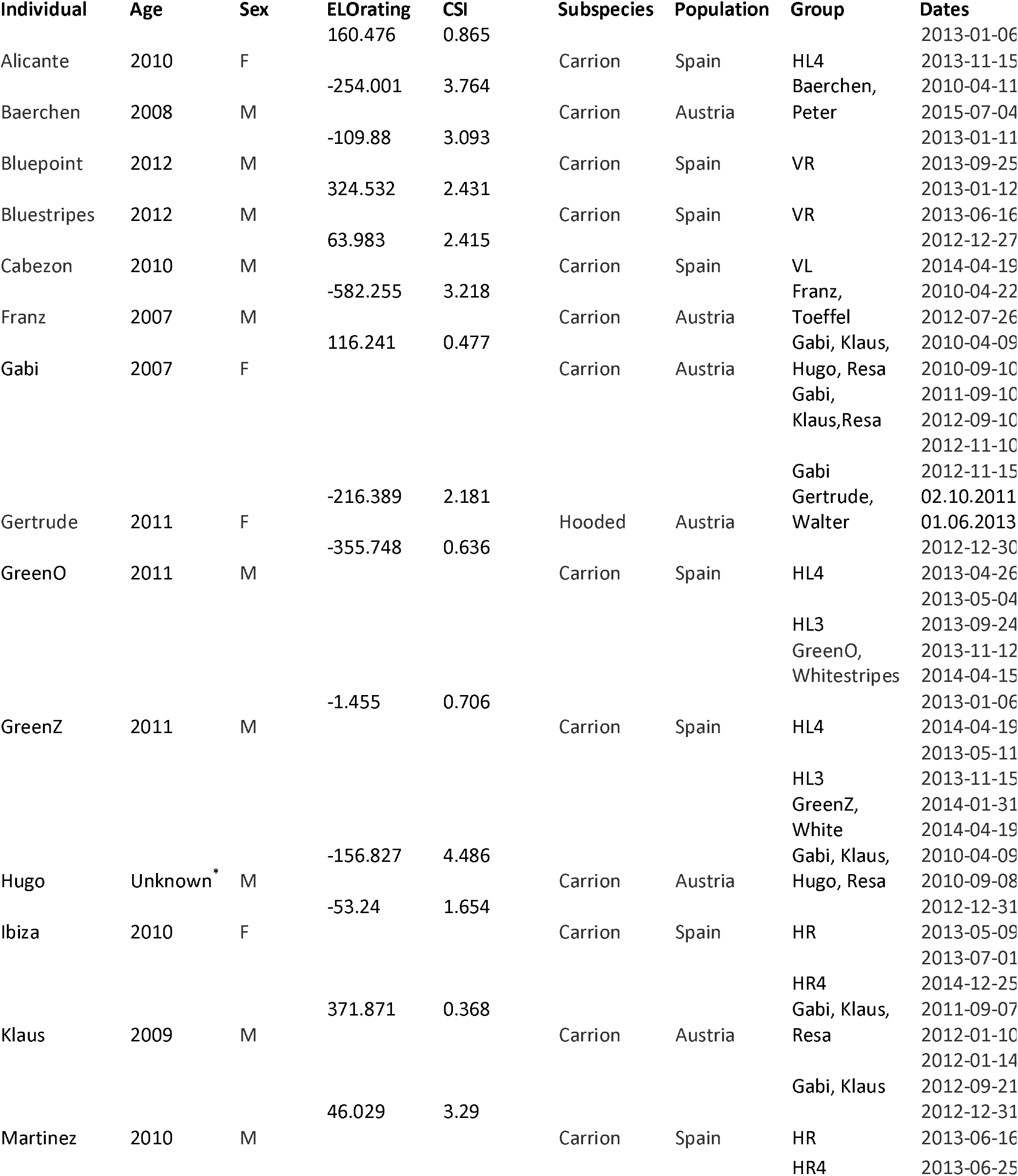
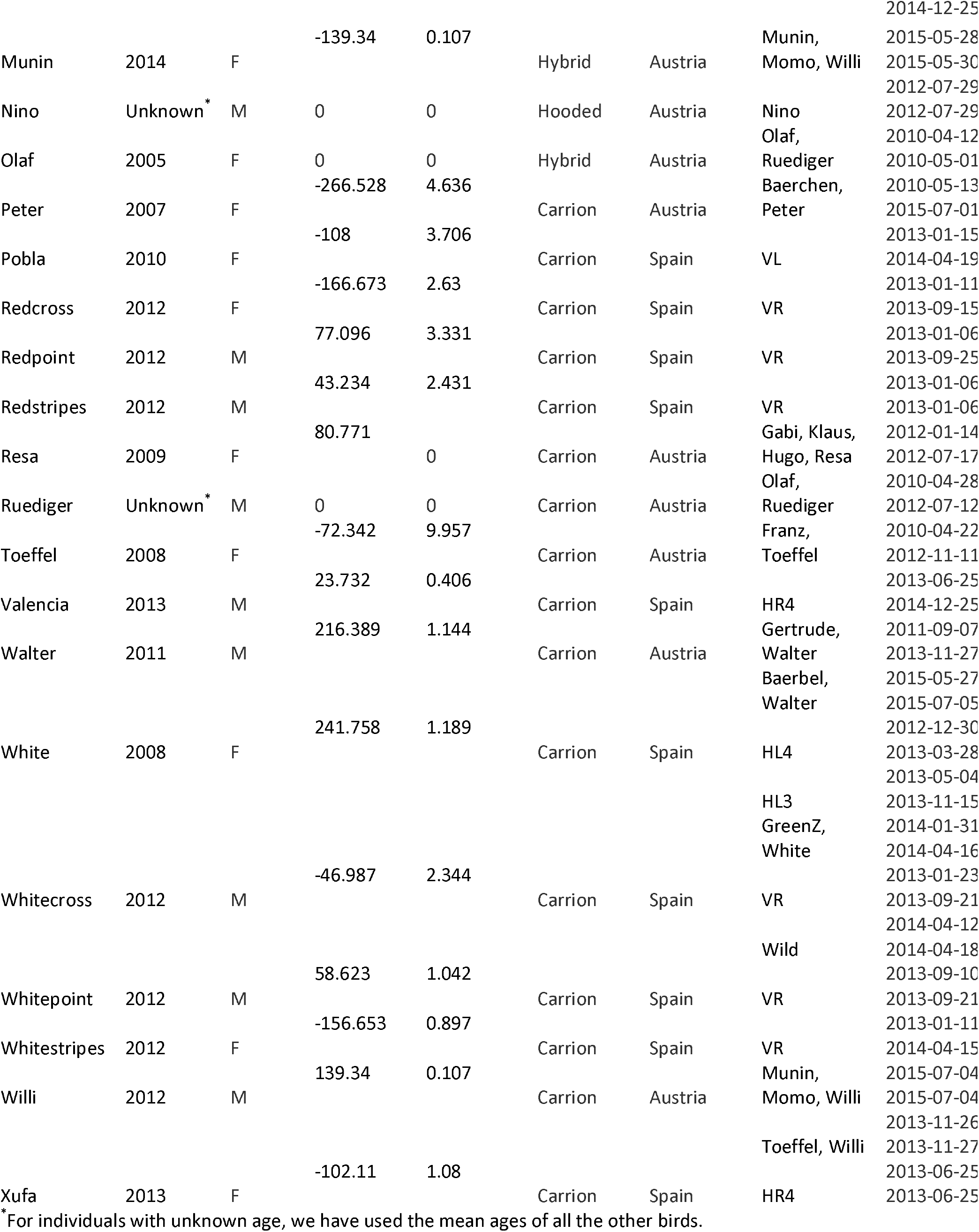
Study subjects. Year of hatching, sex, species, population and group compositions of all individuals contributing to the present study. Dates refer to the times recordings were made in a specific group composition and the number of calls and number of sequences recorded.

### Recording vocalisations and data analysis

Vocalisations have been recorded during regular behavioural observations using either a Sennheiser ME67 directional microphone on a K6 module, and a 13 Marantz PMD661 digital recorder or a SM1 Wildlife acoustics SongMeter. Recordings were opportunistically taken when an observer was present in the enclosure and hence recording durations differ between individuals (Table 1). Crows were generally habituated to the human observer. The human observer identified the calling individual by speaking on the recording.

Onsets and offsets of individual calls have been annotated by CAFW using Audacity® or Raven Pro (K. Lisa Yang Center for Conservation Bioacoustics at the Cornell Lab of Ornithology, 2024). Vocalizations from unknown callers, overlapping vocalizations, incomplete sequences, and single call sequences have been excluded for the present study. Call sequences can be intuitively recognized by human observers, however for the present study we have classified calls with more than one second inter-call duration as separate sequences. This was manually validated by CAFW. In total, we recorded 12,123 calls in 3,726 complete sequences. For details about calls and sequences per individual, see Table 1.

### Statistical analysis

All models were fit using the lme4 (v1.1-35.1) (Bates et al., 2015) package in R (v4.3.1) (R Core Team, 2019). To avoid the many problems associated with p-values, we report mean estimates and 95% Wald confidence intervals and interpret intervals that do not overlap zero as indicating a strong effect. All reported models were manually checked for convergence.

We focus on the Menzerath-Altmann law—a precise and more robust mathematical form of Menzerath’s law (Altmann, 1980; Torre et al., 2021). Here is the standard form of the Menzerath-Altmann law where y is the duration of elements within a sequence composed of x elements, and a, b, and c are parameters controlling the shape of the relationship.

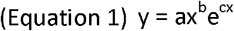

If we set c to 0 and apply some simple algebra, we get a linear model that is the most common form of the law in contemporary linguistics (Hou et al., 2017).

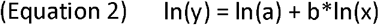

We will follow other studies of the Menzerath-Altmann law in nonhuman animals and use the above linear model (Clink & Lau, 2020; Favaro et al., 2020; Gustison et al., 2016, p. 201; James et al., 2021; Lewis et al., 2023; Stepanov et al., 2023; Vradi, 2021; Youngblood, 2024, 2025). y is usually the mean duration of elements within sequences, but we will use the full distribution of call durations within sequences (Youngblood, 2024; Youngblood; 2025) to avoid spurious “regression to the mean” effects (Ferrer-i-Cancho et al., 2013; Gustison et al., 2016; Milička, 2023) and better capture uncertainty in the models (Youngblood, 2024). We also follow other work in excluding single-element sequences (i.e., with a length of one) from the analysis, which have been shown to depart from Menzerath’s law (G. Torre et al., 2021; Heesen et al., 2019; Hernández-Fernández et al., 2019; Torre et al., 2019).

The “simple” model, used to assess Menzerath’s law separately in carrion crows, hooded crows, and hybrids of carrion and hooded crows, had the following specification:

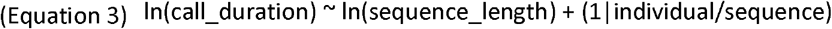

Nested varying intercepts for sequences and individuals are included to account for the repeated measurement of call durations within sequences and for individual variation in call durations. It is also worth noting that we applied the simple model separately to each species, rather than analysing them together with species as a fixed effect to assess species differences. In this case, given the dramatic disparity in sample sizes between the three species, we think it is unwise to make conclusions about the relative strength of Menzerath’s law, and focus on the presence of it instead.

The “complex” model, used to assess individual variation in Menzerath’s law based on demographic factors, had three potential specifications:

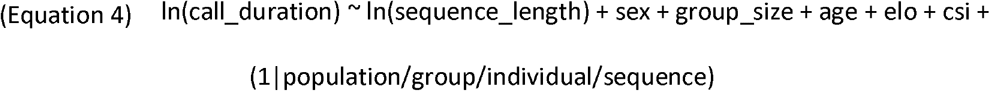

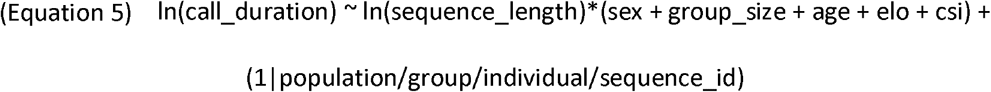

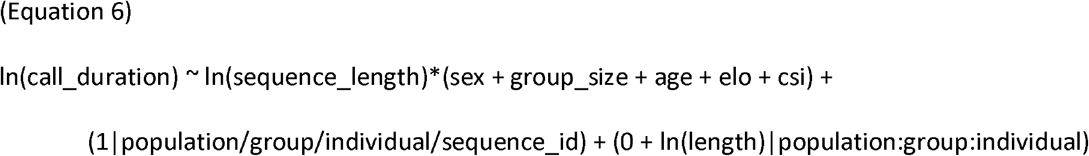

The first of these models includes fixed effects for sequence length, sex, group size, age, ELO, and CSI, as well as nested varying intercepts for population, group, individual, and sequence. The second of these adds interactions between the fixed effect for sequence length and sex, group size, age, ELO, and CSI. The third of these adds additional nested varying slopes that allow for the strength of the effect of sequence length on call duration to vary by population, group, and individual. All three of these models were fit to the data, and the one with the lowest AIC is reported in the results.

Importantly, the complex models were only fit to the data from carrion crows. As seen in Table 1, the data from hooded and hybrid crows are too sparse to assess the effect of demographic factors on Menzerath’s law. For example, both hooded crows come from the Austria population, and both hybrid crows are female, making it impossible to disentangle those factors from species differences. Analysis code and data are available at https://masonyoungblood.github.io/crow_efficiency/.

## Results

In all three subspecies, there is a strong negative relationship between sequence length and call duration, consistent with Menzerath’s law (Figure 1 and Table 2). As mentioned in the methods, we chose to not formally analyse species differences in Menzerath’s law because of the large disparities in sample size, but it is interesting to note that the strength of the law in hybrid crows is situated between carrion and hooded crows (Figure 1 and Table 2).

**Table 2.** The results of the simple model of Menzerath’s law applied separately to call sequences from carrion crows, hooded crows, and hybrids of carrion and hooded crows. 2.5% and 97.5% mark the bounds of the 95% confidence intervals around the point estimates, where intervals that do not overlap zero (i.e., strong effects) are marked with an asterisk.

| Species | Parameter | Estimate | 2.5% (Lower) | 97.5% (Upper) |  |
| --- | --- | --- | --- | --- | --- |
| Carrion Crows | Intercept | 0.018 | -0.161 | 0.197 |  |
|  | Length | -0.533 | -0.559 | -0.507 | * |
| Hooded Crows | Intercept | -0.034 | -1.609 | 1.542 |  |
|  | Length | -0.195 | -0.363 | -0.027 | * |
| Hybrid Crows | Intercept | -0.218 | -0.930 | 0.494 |  |
|  | Length | -0.409 | -0.637 | -0.182 | * |

**Figure 1.**
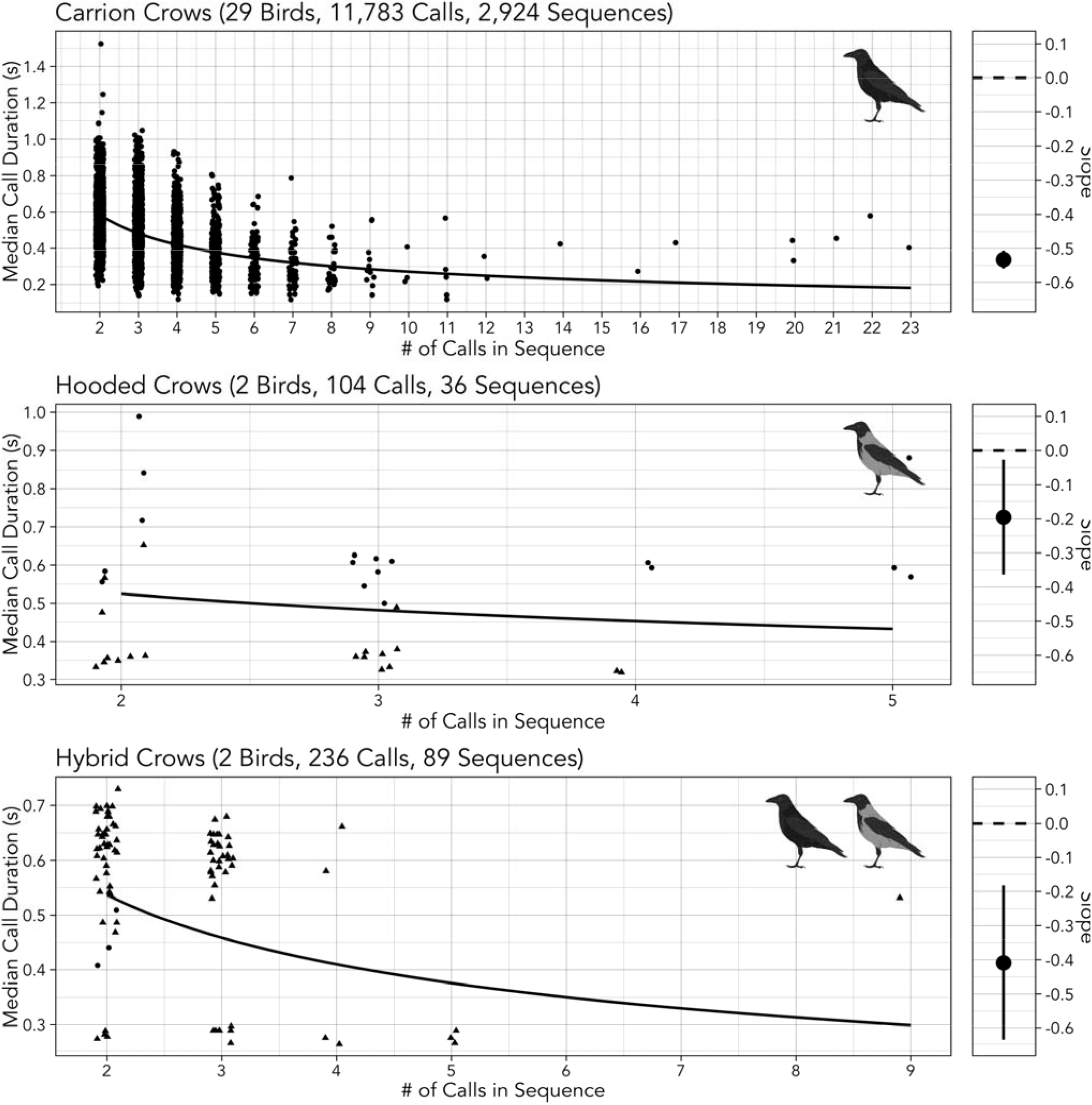
The distribution of median call durations and call sequence lengths (left) and the slope of Menzerath’s law (right) for carrion crows (top), hooded crows (middle), and hybrids of carrion and hooded crows (bottom). Different individuals in hooded crows and hybrid crows are plotted in different symbols. The bars in the slope plot (right) mark the 95% confidence intervals around the point estimates.

**Figure 2.**
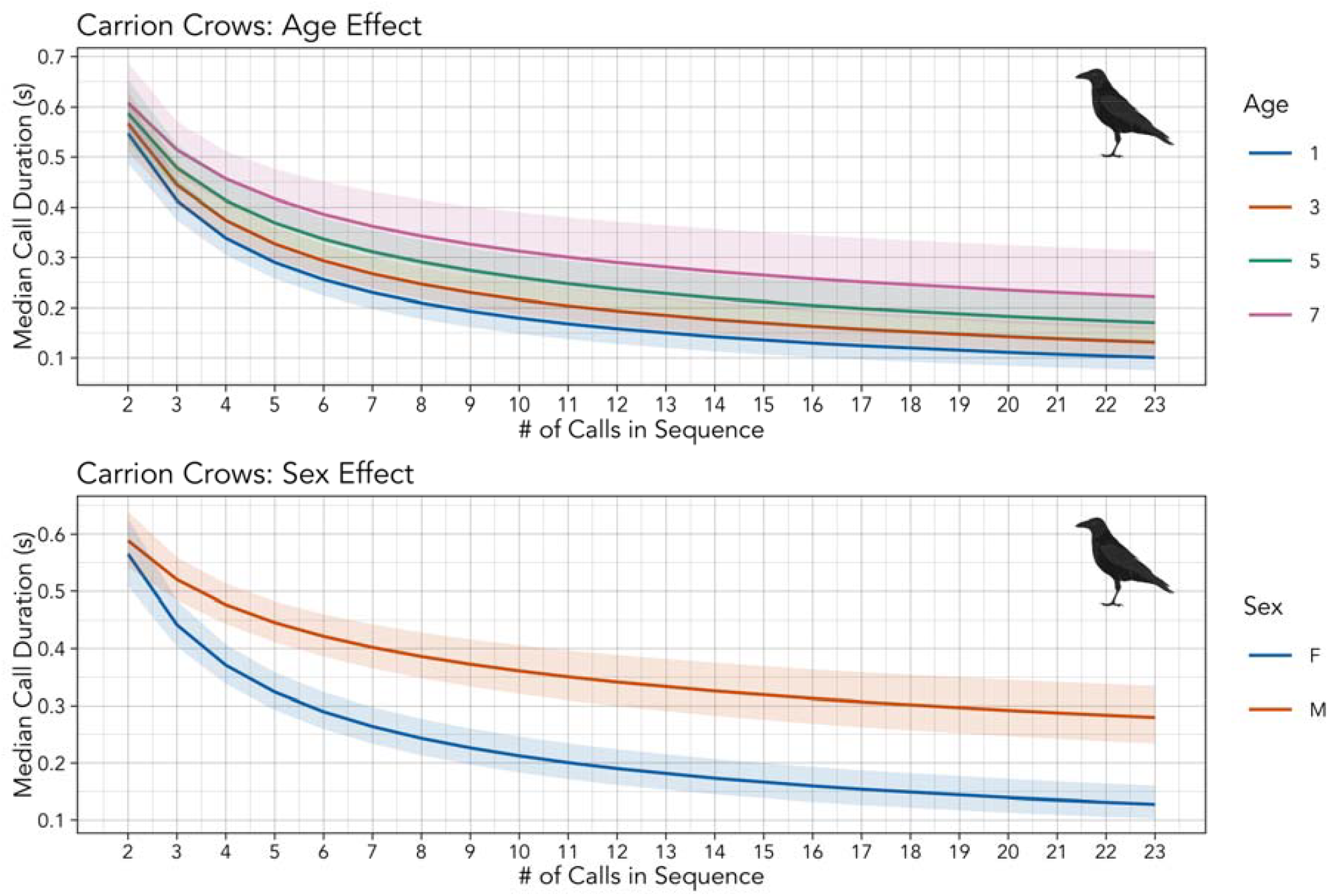
The predicted relationship between element durations and sequence lengths for different levels of age (top) and sex (bottom), from the fitted version of INSERT MODEL.

Of the three complex models fit to carrion crow calls, the third—with interactions between demographic factors and the effect of sequence length, and with varying slopes for individual identity—fit the data best (ΔAIC > 2). Likelihood ratio tests indicate Equation 6 is the best fitting model of individual variation-Equation 5 outcompetes Equation 4 (p = 4.5 x 10^-32), and Equation 6 outcompetes Equation 5 (p = 1.8 x 10^-9). The fact that the third complex model (AIC = 14,653) fits better than the second complex model (AIC = 14,698), which excludes varying slopes for individual identity, suggests that there is meaningful individual variation in Menzerath’s law, even after accounting for demographic factors. The results of the third complex model indicate that demographic factors influence both call duration and the strength of Menzerath’s law (Table 3). Call durations are longer in males, older birds, and more dominant birds, and Menzerath’s law appears to be weaker in both males and older birds (Figure 2).

**Table 3.** The results of the complex model of Menzerath’s law, built to assess individual variation based on demographic factors, applied only to call sequences from carrion crows. 2.5% and 97.5% mark the bounds of the 95% confidence intervals around the point estimates, where intervals that do not overlap zero (i.e., strong effects) are marked with an asterisk. Interactions between demographic factors and Menzerath’s law are marked with colons (e.g., “Length : Age” is the interaction between age and effect of sequence length on call duration).

| Parameter | Estimate | 2.5% (Lower) | 97.5% (Upper) |  |
| --- | --- | --- | --- | --- |
| Intercept | -0.267 | -0.494 | -0.040 | * |
| Length | -0.678 | -0.796 | -0.560 | * |
| Sex (M vs. F) | 0.541 | 0.254 | 0.827 | * |
| Group Size | -0.107 | -0.214 | 0.001 |  |
| Age | 0.142 | 0.086 | 0.198 | * |
| ELO | 0.201 | 0.078 | 0.324 | * |
| CSI | 0.081 | -0.087 | 0.250 |  |
| Length : Sex (M vs. F) | 0.338 | 0.183 | 0.494 | * |
| Length : Group Size | -0.043 | -0.108 | 0.022 |  |
| Length : Age | 0.066 | 0.014 | 0.119 | * |
| Length : ELO | 0.045 | -0.019 | 0.109 |  |
| Length : CSI | -0.033 | -0.114 | 0.048 |  |

## Discussion

In the present study we show that the call sequences of crows—specifically carrion crows, hooded crows, and hybrids of carrion and hooded crows—adhere to Menzerath’s law. In other words, call sequences with more calls are composed of shorter calls, which is suggested to lead to greater communicative efficiency in human language (Piantadosi et al., 2011). To our knowledge, this is the first time efficiency laws have been shown in vocal communication of corvids, adding to the widespread evidence for Menzerath’s law in both human and non-human communication systems (Deng et al., 2024; Gustison et al., 2016; Semple et al., 2010; Youngblood, 2024, 2024, 2025). Interestingly, within carrion crows, adherence to Menzerath’s Law was modulated by sex and age. Males’ calls are longer, and their adherence to Menzerath’s law is weaker, which may reflect sex-specific roles in social organization or differences in vocal usage patterns. Linguistic laws previously mostly have been investigated in song, which in many species is sexually selected in males to attract females (Clink & Lau, 2020; James et al., 2021; Valente et al., 2021; Youngblood, 2024, 2025). Calls in corvids often function as contact calls (Anjos & Vielliard, 1993; Berger & Ligon, 1977; McCaig et al., 2015; Roskaft & Espmark, 1982; Sitasuwan & Thaler, 1985), food calls (Bugnyar et al., 2001; Mates et al., 2015; Roehmholdt, 2019; Roskaft & Espmark, 1982; Szipl & Bugnyar, 2014), territory calls (Mates et al., 2015; Sabol et al., 2022; Tanimoto et al., 2017), or alert signals (Chamberlain & Cornwell, 1971; Conner, 1985; Cornell et al., 2012; Ellis, 2009; Mates et al., 2015; Pendergraft & Marzluff, 2019; Tanimoto et al., 2017; Yorzinski & Vehrencamp, 2009) and are typically emitted by both sexes. In rooks, males produced call units with lower diversity and gradation than females and individual males produced different call repertoires, whereas females produced more similar call repertoires to each other (Martin et al., 2024). Our study indicates that female carrion crows might communicate more efficiently compared to males. This is different to a previous study in lemurs, *Indri indri*, which showed no sex difference in adherence to Menzerath’s law (Valente et al., 2021).

Surprisingly, younger individuals displayed stronger adherence to Menzerath’s Law compared to older crows. This finding contrasts with our original prediction, that efficiency in vocal production would increase with age. This finding contrasts previous findings in zebra finches, *Taeniopygia castanotis*, were song elements and gaps shorten over development (Glaze & Troyer, 2013) and humans, were vocal efficiency increases with age (Tang & Stathopoulos, 1995). Songbirds reared without auditory experiences that guide vocal development still adhere to Menzerath’s law, indicating that vocal efficiency is linked to vocal production mechanisms rather than learned behaviour (James et al., 2021). Our results could indicate that carrion crow vocalisations are limited to vocal efficiency at younger age and become more flexible with increased age. Since call duration overall also increases with age in our analysis, we suggest the need for informativeness of calls may outweigh the importance of efficiency as crows age. Further studies on the ontogeny of adherence to Menzerath’s law are important.

Group size, dominance status, and the strength of affiliative relationships did not significantly influence adherence to Menzerath’s Law, although call duration itself was longer in more dominant birds. This suggests that, in carrion crows, the structural efficiency of vocal sequences may be relatively stable across different social contexts. However, it should be noted that our data stems from captive individuals, where social context such as group composition were under human control, hence the full range of social contexts might not be captured in the data. Overall, our findings highlight the presence of Menzerath’s Law in carrion crow communication and point to individual-level social factors, rather than broader group dynamics, as influencing the degree of adherence. Future research should explore whether these patterns hold across different communicative contexts such as different call-types and in other corvid species, to better understand the interplay between social structure, cognitive demands, and communicative efficiency.

## References

Altmann, G. (1980). Prolegomena to Menzerath’s law. Glottometrika, 2(2), 1–10.

Anjos, L. D., & Vielliard, J. M. E. (1993). Repertoire of the acoustic communication of the azure jay Cyanocorax caeruleus (Vieillot) (Aves, Corvidae). Revista Brasileira de Zoologia, 10(4), 657–664. 10.1590/S0101-81751993000400011

Archie, E. A., Tung, J., Clark, M., Altmann, J., & Alberts, S. C. (2014). Social affiliation matters: Both same-sex and opposite-sex relationships predict survival in wild female baboons. Proceedings of the Royal Society B: Biological Sciences, 281(1793), 20141261. 10.1098/rspb.2014.1261

AviList Core Team. (2025). AviList: The Global Avian Checklist, v2025. [Dataset]. 10.2173/avilist.v2025

Bates, D., Mächler, M., Bolker, B., & Walker, S. (2015). Fitting linear mixed-effects models using lme4. Journal of Statistical Software, 67(1), 1–48.

Berger, L. R., & Ligon, J. D. (1977). Vocal communication and individual recognition in the pinon jay, Gymnorhinus cyanocephalus. Animal Behaviour, 25, 567–584. 10.1016/0003-3472(77)90107-5

Bezerra, B. M., Souto, A. S., Radford, A. N., & Jones, G. (2011). Brevity is not always a virtue in primate communication. Biology Letters, 7(1), 23–25. 10.1098/rsbl.2010.0455

Brecht, K. F., Hage, S. R., Gavrilov, N., & Nieder, A. (2019). Volitional control of vocalizations in corvid songbirds. PLOS Biology, 17(8), e3000375. 10.1371/journal.pbio.3000375

Brenowitz, E. A., Margoliash, D., & Nordeen, K. W. (1997). An introduction to birdsong and the avian song system. Journal of Neurobiology, 33(5), 495–500. 10.1002/(SICI)1097-4695(19971105)33:5<495::AID-NEU1>3.0.CO;2-#

Bugnyar, T., Kijne, M., & Kotrschal, K. (2001). Food calling in ravens: Are yells referential signals? Animal Behaviour, 61(5), 949–958. 10.1006/anbe.2000.1668

Chamberlain, D. R., & Cornwell, G. W. (1971). Selected vocalizations of the common crow. The Auk, 88, 613–634.

Choi, N., Adams, M., Fowler-Finn, K., Knowlton, E., Rosenthal, M., Rundus, A., Santer, R. D., Wilgers, D., & Hebets, E. A. (2022). Increased signal complexity is associated with increased mating success. Biology Letters, 18(5), 20220052. 10.1098/rsbl.2022.0052

Clink, D. J., & Lau, A. R. (2020). Adherence to Menzerath’s Law is the exception (not the rule) in three duetting primate species. Royal Society Open Science, 7(11), 201557. 10.1098/rsos.201557

Conner, R. N. (1985). Vocalizations of common ravens in Virginia. The Condor, 87(3), 379– 388. 10.2307/1367219

Cornell, H. N., Marzluff, J. M., & Pecoraro, S. (2012). Social learning spreads knowledge about dangerous humans among American crows. Proceedings of the Royal Society B: Biological Sciences, 279(1728), 499–508. 10.1098/rspb.2011.0957

Demartsev, V., Gordon, N., Barocas, A., Bar-Ziv, E., Ilany, T., Goll, Y., Ilany, A., & Geffen, E. (2019). The “Law of Brevity” in animal communication: Sex-specific signaling optimization is determined by call amplitude rather than duration. Evolution Letters, 3(6), 623–634. 10.1002/evl3.147

Deng, K., He, Y.-X., Wang, X.-P., Wang, T.-L., Wang, J.-C., Chen, Y.-H., & Cui, J.-G. (2024). Hainan frilled treefrogs’ calls partially conform to Menzerath–Altmann’s law, but oppose Zipf’s law of abbreviation. Animal Behaviour, 213, 51–59. 10.1016/j.anbehav.2024.04.011

Ellis, J. M. S. (2009). Anti[predator signals as advertisements: Evidence in white[throated magpie[jays. Ethology, 115(6), 522–532. 10.1111/j.1439-0310.2009.01631.x

Endler, J. A. (1993). Some general comments on the evolution and design of animal communication systems. Philosophical Transactions: Biological Sciences, 340(1292), 215–225.

Favaro, L., Gamba, M., Cresta, E., Fumagalli, E., Bandoli, F., Pilenga, C., Isaja, V., Mathevon, N., & Reby, D. (2020). Do penguins’ vocal sequences conform to linguistic laws? Biology Letters, 16(2), 20190589. 10.1098/rsbl.2019.0589

Ferrer□i□Cancho, R., Hernández[Fernández, A., Lusseau, D., Agoramoorthy, G., Hsu, M. J., & Semple, S. (2013). Compression as a universal principle of animal behavior. Cognitive Science, 37(8), 1565–1578. 10.1111/cogs.12061

Freeberg, T. M., & Krams, I. (2015). Does social complexity link vocal complexity and cooperation? Journal of Ornithology, 156(S1), 125–132. 10.1007/s10336-015-1233-2

G. Torre, I., Dębowski, Ł., & Hernández-Fernández, A. (2021). Can Menzerath’s law be a criterion of complexity in communication? PLOS ONE, 16(8), e0256133. 10.1371/journal.pone.0256133

Gibson, E., Futrell, R., Piantadosi, S. P., Dautriche, I., Mahowald, K., Bergen, L., & Levy, R. (2019). How efficiency shapes human language. Trends in Cognitive Sciences, 23(5), 389–407. 10.1016/j.tics.2019.02.003

Glaze, C. M., & Troyer, T. W. (2013). Development of temporal structure in zebra finch song. Journal of Neurophysiology, 109(4), 1025–1035. 10.1152/jn.00578.2012

Gustison, M. L., Semple, S., Ferrer-i-Cancho, R., & Bergman, T. J. (2016). Gelada vocal sequences follow Menzerath’s linguistic law. Proceedings of the National Academy of Sciences, 113(19). 10.1073/pnas.1522072113

Heesen, R., Hobaiter, C., Ferrer-i-Cancho, R., & Semple, S. (2019). Linguistic laws in chimpanzee gestural communication. Proceedings of the Royal Society B: Biological Sciences, 286(1896), 20182900. 10.1098/rspb.2018.2900

Hernández-Fernández, A., G. Torre, I., Garrido, J.-M., & Lacasa, L. (2019). Linguistic Laws in speech: The case of Catalan and Spanish. Entropy, 21(12), 1153. 10.3390/e21121153

Hou, R., Huang, C.-R., Do, H. S., & Liu, H. (2017). A study on correlation between Chinese sentence and constituting clauses based on the Menzerath-Altmann law. Journal of Quantitative Linguistics, 24(4), 350–366. 10.1080/09296174.2017.1314411

Huang, M., Ma, H., Ma, C., Garber, P. A., & Fan, P. (2020). Male gibbon loud morning calls conform to Zipf’s law of brevity and Menzerath’s law: Insights into the origin of human language. Animal Behaviour, 160, 145–155. 10.1016/j.anbehav.2019.11.017

Igamberdiev, A. U. (2023). Overcoming the limits of natural computation in biological evolution toward the maximization of system efficiency. Biological Journal of the Linnean Society, 139(4), 539–554. 10.1093/biolinnean/blac093

James, L. S., Mori, C., Wada, K., & Sakata, J. T. (2021). Phylogeny and mechanisms of shared hierarchical patterns in birdsong. Current Biology, 31(13), 2796-2808.e9. 10.1016/j.cub.2021.04.015

K. Lisa Yang Center for Conservation Bioacoustics at the Cornell Lab of Ornithology. (2024). Raven Pro: Interactive Sound Analysis Software [Computer software]. (Version 1.6.5) [Computer software]. https://www.ravensoundsoftware.com/

Krams, I., Krama, T., Freeberg, T. M., Kullberg, C., & Lucas, J. R. (2012). Linking social complexity and vocal complexity: A parid perspective. Philosophical Transactions of the Royal Society B: Biological Sciences, 367(1597), 1879–1891. 10.1098/rstb.2011.0222

Lewis, R. N., Kwong, A., Soma, M., De Kort, S. R., & Gilman, R. T. (2023). Java sparrow song conforms to Menzerath’s Law but not Zipf’s Law of Abbreviation. 10.1101/2023.12.13.571437

Liao, D. A., Brecht, K. F., Veit, L., & Nieder, A. (2024). Crows “count” the number of self-generated vocalizations. Science, 384(6698), 874–877. 10.1126/science.adl0984

Luef, E. M., Ter Maat, A., & Pika, S. (2017). Vocal similarity in long-distance and short-distance vocalizations in raven pairs ( Corvus corax ) in captivity. Behavioural Processes, 142, 1–7. 10.1016/j.beproc.2017.05.013

Manser, M. B., Jansen, D. A. W. A. M., Graw, B., Hollén, L. I., Bousquet, C. A. H., Furrer, R. D., & Le Roux, A. (2014). Vocal Complexity in Meerkats and Other Mongoose Species. In Advances in the Study of Behavior (Vol. 46, pp. 281–310). Elsevier. 10.1016/B978-0-12-800286-5.00006-7

Martin, K., Cornero, F. M., Clayton, N. S., Adam, O., Obin, N., & Dufour, V. (2024). Vocal complexity in a socially complex corvid: Gradation, diversity and lack of common call repertoire in male rooks. Royal Society Open Science, 11(1), 231713. 10.1098/rsos.231713

Mates, E. A., Tarter, R. R., Ha, J. C., Clark, A. B., & McGowan, K. J. (2015). Acoustic profiling in a complexly social species, the American crow: Caws encode information on caller sex, identity and behavioural context. Bioacoustics, 24(1), 63–80. 10.1080/09524622.2014.933446

McCaig, T., Brown, M., & Jones, D. N. (2015). Exploring possible functions of vocalisations in the Torresian Crow Corvus orru. Australian Field Ornithology, 32, 201–208.

Menzerath, P. (1954). Die Architektonik des deutschen Wortschatzes. Dümmler.

Milička, J. (2023). Menzerath’s law: Is it just regression toward the mean? Glottometrics, 55, 1– 16. 10.53482/2023_55_409

Neumann, C., Duboscq, J., Dubuc, C., Ginting, A., Irwan, A. M., Agil, M., Widdig, A., & Engelhardt, A. (2011). Assessing dominance hierarchies: Validation and advantages of progressive evaluation with Elo-rating. Animal Behaviour, 82(4), 911–921. 10.1016/j.anbehav.2011.07.016

Peckre, L., Kappeler, P. M., & Fichtel, C. (2019). Clarifying and expanding the social complexity hypothesis for communicative complexity. Behavioral Ecology and Sociobiology, 73(1), 11. 10.1007/s00265-018-2605-4

Pendergraft, L. T., & Marzluff, J. M. (2019). Fussing over food: Factors affecting the vocalizations American crows utter around food. Animal Behaviour, 150, 39–57. 10.1016/j.anbehav.2019.01.024

Piantadosi, S. T., Tily, H., & Gibson, E. (2011). Word lengths are optimized for efficient communication. Proceedings of the National Academy of Sciences, 108(9), 3526–3529. 10.1073/pnas.1012551108

Pollard, K. A., & Blumstein, D. T. (2012). Evolving communicative complexity: Insights from rodents and beyond. Philosophical Transactions of the Royal Society B: Biological Sciences, 367(1597), 1869–1878. 10.1098/rstb.2011.0221

R Core Team. (2019). R: a language and environment for statistical computing. (Version 4.3.1.) [Computer software]. http://www.r723project.org/

Risueno-Segovia, C., Dohmen, D., Gultekin, Y. B., Pomberger, T., & Hage, S. R. (2023). Linguistic law-like compression strategies emerge to maximize coding efficiency in marmoset vocal communication. Proceedings of the Royal Society B: Biological Sciences, 290(2007), 20231503. 10.1098/rspb.2023.1503

Roehmholdt, C. E. (2019). Using vocalizations to determine the capacity for future cognition in the common raven (Corvus Corax) [Doctoral Thesis]. Evergreen State College.

Roskaft, E., & Espmark, Y. (1982). Vocal communication by the rook Corvus frugilegus during the breeding season. Ornis Scandinavica, 13(1), 38. 10.2307/3675971

Sabol, A. C., Greggor, A. L., Masuda, B., & Swaisgood, R. R. (2022). Testing the maintenance of natural responses to survival-relevant calls in the conservation breeding population of a critically endangered corvid (Corvus hawaiiensis). Behavioral Ecology and Sociobiology, 76(1), 21. 10.1007/s00265-022-03130-8

Safryghin, A., Cross, C., Fallon, B., Heesen, R., Ferrer-i-Cancho, R., & Hobaiter, C. (2022). Variable expression of linguistic laws in ape gesture: A case study from chimpanzee sexual solicitation. Royal Society Open Science, 9(11), 220849. 10.1098/rsos.220849

Sánchez-Tójar, A., Schroeder, J., & Farine, D. R. (2018). A practical guide for inferring reliable dominance hierarchies and estimating their uncertainty. Journal of Animal Ecology, 87(3), 594–608. 10.1111/1365-2656.12776

Sandoval, L., & Graham, B. (2025). Songs and calls: Perspectives on creating a global definition. Ornitología Neotropical, 35(2). 10.58843/ornneo.v35i2.1361

Semple, S., Ferrer-i-Cancho, R., & Gustison, M. L. (2022). Linguistic laws in biology. Trends in Ecology & Evolution, 37(1), 53–66. 10.1016/j.tree.2021.08.012

Semple, S., Hsu, M. J., & Agoramoorthy, G. (2010). Efficiency of coding in macaque vocal communication. Biology Letters, 6(4), 469–471. 10.1098/rsbl.2009.1062

Silk, J. B., Beehner, J. C., Bergman, T. J., Crockford, C., Engh, A. L., Moscovice, L. R., Wittig, R. M., Seyfarth, R. M., & Cheney, D. L. (2010). Female chacma baboons form strong, equitable, and enduring social bonds. Behavioral Ecology and Sociobiology, 64(11), 1733–1747. 10.1007/s00265-010-0986-0

Sitasuwan, N., & Thaler, E. (1985). Vocal inventory and communication in the Chough (Pyrrhocorax pyrrhocorax), in the Alpine Chough (P. graculus) and in their hybrids. Journal für Ornithologie, 126, 181–193.

Stepanov, A., Zhivomirov, H., Nedelchev, I., & Stateva, P. (2023). Bottlenose dolphins’ broadband clicks are structured for communication. 10.1101/2023.01.11.523588

Suzuki, T. N. (2014). Communication about predator type by a bird using discrete, graded and combinatorial variation in alarm calls. Animal Behaviour, 87, 59–65. 10.1016/j.anbehav.2013.10.009

Szipl, G., & Bugnyar, T. (2014). Craving ravens: Individual ‘haa’ call rates at feeding sites as cues to personality and levels of fission-fusion dynamics? Animal Behavior and Cognition, 1(3), 265. 10.12966/abc.08.04.2014

Szipl, G., Ringler, E., & Bugnyar, T. (2018). Attacked ravens flexibly adjust signalling behaviour according to audience composition. Proceedings of the Royal Society B: Biological Sciences, 285(1880), 20180375. 10.1098/rspb.2018.0375

Tang, J., & Stathopoulos, E. T. (1995). Vocal efficiency as a function of vocal intensity: A study of children, women, and men. The Journal of the Acoustical Society of America, 97(3), 1885–1892. 10.1121/1.412062

Tanimoto, A. M., Hart, P. J., Pack, A. A., & Switzer, R. (2017). Vocal repertoire and signal characteristics of ‘Alalā, the Hawaiian Crow ( Corvus hawaiiensis ). The Wilson Journal of Ornithology, 129(1), 25–35. 10.1676/1559-4491-129.1.25

Torre, I. G., Luque, B., Lacasa, L., Kello, C. T., & Hernández-Fernández, A. (2019). On the physical origin of linguistic laws and lognormality in speech. Royal Society Open Science, 6(8), 191023. 10.1098/rsos.191023

Valente, D., De Gregorio, C., Favaro, L., Friard, O., Miaretsoa, L., Raimondi, T., Ratsimbazafy, J., Torti, V., Zanoli, A., Giacoma, C., & Gamba, M. (2021). Linguistic laws of brevity: Conformity in Indri indri. Animal Cognition, 24(4), 897–906. 10.1007/s10071-021-01495-3

Vradi, A. A. (2021). Dolphin communication: A quantitative linguistics approach [Master’s thesis]. Universitat Politècnica de Catalunya.

Wascher, C. A. F. (2021). Association between social factors and gastrointestinal parasite product excretion in a group of non-cooperatively breeding carrion crows. Behavioral Ecology and Sociobiology, 75(2), 30. 10.1007/s00265-021-02967-9

Wascher, C. A. F., Canestrari, D., & Baglione, V. (2019). Affiliative social relationships and coccidian oocyst excretion in a cooperatively breeding bird species. Animal Behaviour, 158, 121–130. 10.1016/j.anbehav.2019.10.009

Wascher, C. A. F., & Reynolds, S. (2025). Vocal communication in corvids: A systematic review. Animal Behaviour, 221, 123073. 10.1016/j.anbehav.2024.123073

Wascher, C. A. F., Waterhouse, G., & Beheim, B. A. (2025). Vocal mimicry in corvids. BioRxiv. 10.1101/2025.03.26.645457

Yorzinski, J. L., & Vehrencamp, S. L. (2009). The effect of predator type and danger level on the mob calls of the American crow. The Condor, 111(1), 159–168. 10.1525/cond.2009.080057

Youngblood, M. (2024). Language-like efficiency and structure in house finch song. Proceedings of the Royal Society B: Biological Sciences, 291(2020), 20240250. 10.1098/rspb.2024.0250

Youngblood, M. (2025). Language-like efficiency in whale communication. Science Advances, 11, eads6014. 10.31234/osf.io/tduab

Zhang, C., Zheng, Z., Lucas, J. R., Wang, Y., Fan, X., Zhao, X., Feng, J., Sun, C., & Jiang, T. (2024). Do bats’ social vocalizations conform to Zipf’s law and the Menzerath-Altmann law? iScience, 27(7), 110401. 10.1016/j.isci.2024.110401

Zipf, G. K. (1949). Human behavior and the principle of least effort: An introduction to human ecology. Addison-Wesley Press.

